# A fluorescent sensor for the real-time measurement of extracellular oxytocin dynamics in the brain

**DOI:** 10.1101/2021.07.30.454450

**Authors:** Daisuke Ino, Hiroshi Hibino, Masaaki Nishiyama

**Affiliations:** Department of Histology and Cell Biology, Graduate School of Medical Sciences, Kanazawa University, Japan; Department of Pharmacology, Graduate School of Medicine, Osaka University, Japan

## Abstract

Oxytocin (OT), a hypothalamic neuropeptide that acts as a neuromodulator in the brain, orchestrates a variety of animal behaviors. However, the relationship between brain OT dynamics and complex animal behaviors remains largely elusive, partly because of the lack of a suitable technique for its real-time recording *in vivo*. Here, we describe MTRIA_OT_, a G protein-coupled receptor-based green fluorescent OT sensor with a large dynamic range, optimal affinity, ligand specificity to OT orthologs, minimal effects on downstream signaling, and long-term fluorescence stability. By combining viral gene delivery and fiber photometry-mediated fluorescence measurements, we demonstrated the utility of MTRIA_OT_ for real-time detection of brain OT dynamics in living mice. Importantly, MTRIA_OT_-mediated measurements revealed “OT oscillation,” a hitherto unknown rhythmic change in OT levels in the brain. MTRIA_OT_ will allow the analysis of OT dynamics in a wide variety of physiological and pathological processes.

## INTRODUCTION

Oxytocin (OT) is a neuropeptide that was initially discovered in the early 20th century as a component of pituitary extracts that induces powerful uterine contractions^1,2^. It has since attracted attention as a key regulator of many physiological processes. In addition to its well-characterized roles as a peripheral hormone that promotes childbirth and lactation, OT is also released as a neuromodulator within the brain, thus participating in the regulation of diverse physiological functions such as sensory processing, feeding control, social cognition, and emotion^3^. To exert these central effects, OT is primarily supplied from neurons in the paraventricular nucleus (PVN) and supraoptic nucleus of the hypothalamus. This OT acts on target brain regions including the limbic regions, sensory cortex, and brainstem, where the OT receptor (OTR), a G protein-coupled receptor (GPCR), is highly expressed^4–6^. The release of OT in the brain is likely evoked by external inputs such as sensory cues, food intake, and social rewards^7–10^; however, it can also be controlled in an autoregulatory manner, potentially by the signaling associated with biological clocks^11–13^. It has been reported that the impairment of OT signaling in the brain may underlie the cognitive and emotional dysfunction associated with neurodevelopmental disorders (e.g., autism spectrum disorders and schizophrenia) and brain aging^14,15^. Nevertheless, despite its importance in both health and disease, the relationship between animal behaviors and the temporal distributions of OT in the brain remains largely elusive.

In addition to its important roles as an endogenous ligand, OT has emerged as a potential therapeutic agent for psychiatric disorders based on a finding that exogenous OT administration enhances positive emotions in humans^16^. Furthermore, OT administered peripherally, such as via intranasal or intravenous routes, was originally believed to reach the brain and exert therapeutic effects. However, the potency of exogenously applied OT on central disorders is currently controversial^17–20^. One of the biggest questions is whether OT administered from a peripheral route can efficiently reach the brain through the nose–brain pathway and/or the blood–brain barrier; this has not yet been well addressed^17^.

In this context, techniques that allow the detection of brain OT dynamics are urgently needed. However, the currently available methods have critical limitations. Although the levels of OT in the brain have classically been inferred from levels in peripheral body fluids, such as plasma, saliva, and urine, peripheral OT levels do not always reflect central OT levels, mainly because of the impermeability of the blood–brain barrier to OT^21^. For the direct measurement of brain OT levels, biochemical analyses of sampled cerebrospinal fluid or dialysate collected through microdialysis probes have been the most prevailing approach^12,22,23^. More recently, a reporter gene-based assay (iTango) has been performed for the detection of OT-stimulated cells in the brain^24^. However, the slow sampling rate of these techniques (usually from several tens of minutes to several hours) is unsuitable for tracking brain OT dynamics in real time.

Optical measurements using genetically encoded fluorescent sensors are a potential technique for the *in vivo* detection of signaling molecules, and have good sensitivity, high specificity, and excellent spatiotemporal resolution^25^. However, the utility of optical measurements hinges on the successful development of sensitive fluorescent sensors for specific ligands. Recently, fluorescent sensors for neurotransmitters and neuromodulators such as dopamine, acetylcholine, norepinephrine, and adenosine have been engineered as chimera proteins, consisting of a GPCR with a fluorescent protein (FP) replacing the amino acids in its third intracellular loop (IL3)^26–30^. These sensors show robust fluorescence responses upon agonist binding, likely because of optimal coupling between the conformational change of the IL3 and the environmental change of the FP chromophore. Because the ligands of GPCRs cover a large proportion of extracellular signaling molecules^31^, GPCRs are expected to be used as the ligand-binding scaffolds in the development of fluorescent sensors for various ligands.

Here, we developed a new green fluorescent OT sensor named MTRIA_OT_, which is composed of a medaka OTR and a circularly permutated green FP (cpGFP)-based fluorescent module named MTRIA (<u>*M*</u>odular fluorescence unit fused with <u>*TR*</u>ansmembrane region-to-<u>*I*</u>ntr<u>*A*</u>cellular loop linkers). We demonstrated that MTRIA_OT_ can be used to analyze changes in brain OT levels following exogenous OT application in anesthetized mice. Moreover, MTRIA_OT_ was able to reveal the existence of oscillatory OT dynamics in a cortical region of freely behaving adult mice, the patterns of which were altered by anesthesia, food deprivation, and aging. Finally, we also demonstrated the utility of MTRIA, the fluorescent module of MTRIA_OT_, for the efficient development of a variety of GPCR-based fluorescent sensors. Together, our findings indicate that MTRIA_OT_ offers opportunities for the detection of OT dynamics in the living brain, and potentially expands the repertoire of GPCR-based fluorescent sensors for extracellular ligands.

## RESULTS

### Development of an ultrasensitive fluorescent OT sensor

To develop a GPCR-based fluorescent sensor for extracellular OT, we screened for an optimal OTR that showed good targeting to the plasma membrane (PM). Because most species have only a single subtype of OTR, we examined the PM localization of six OTRs derived from different groups of vertebrates (human, mouse, chicken, snake, frog, and medaka) in human embryonic kidney 293T (HEK293T) cells. We chose the medaka OTR (meOTR) as the scaffold for the development of the new fluorescent sensor because its localization had the best correlation with that of a PM marker (Fig. 1a–c and Extended Data Fig. 1a).

**Figure 1.**
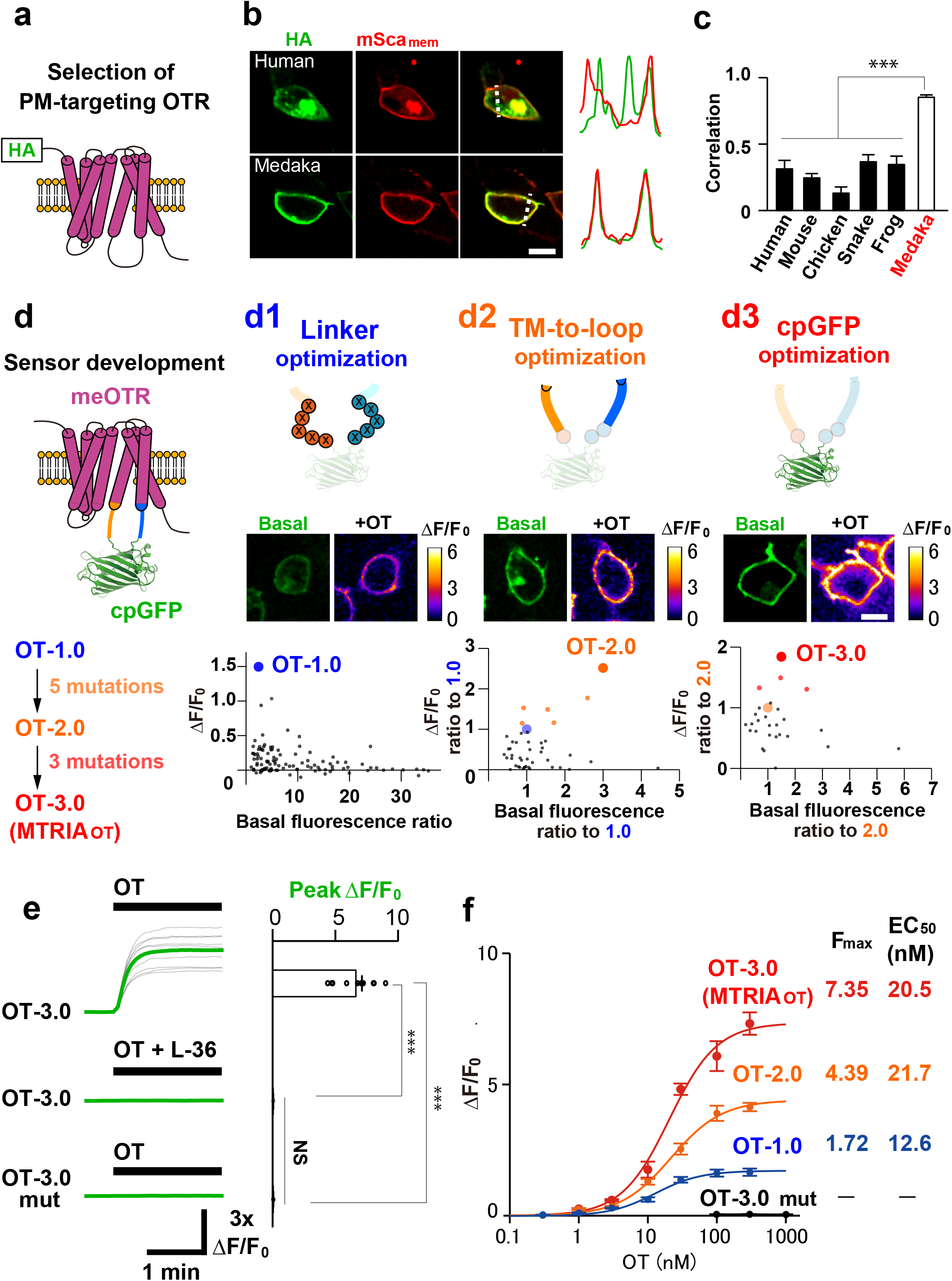
Development of a new fluorescent OT sensor. **a**, Schematic representation of a PM-targeting OTR conjugated with an HA-tag at the N-terminus. **b**, Images of HEK293T cells co-expressing an HA-tagged OTR (HA: green) and a PM-targeted mScarlet (mSca_mem_: red). The right traces compare the normalized fluorescence intensities of HA and mSca_mem_ signals on the dotted lines. **c**, Summary of the Pearson correlation coefficients of fluorescence signals in **b** (*n* = 8, 17, 10, 10, 12, 10, and 17 cells for human, mouse, chicken, snake, frog, and medaka, respectively). Statistics: one-way ANOVA (*F*_5,68_ = 2.35, *P* = 4.9×10^−19^) with Bonferroni *post-hoc* test (*P* = 3.3×10^−9^: human vs. medaka, *P* = 8.9×10^−9^: mouse vs. medaka, *P* = 5.4×10^−14^: chicken vs. medaka, *P* = 2.4×10^−19^: snake vs. medaka, *P* = 2.3×10^−8^: frog vs. medaka). **d**, Development of an ultrasensitive fluorescent OT sensor over three step-screening; optimization of linker regions (**d1**), TM-to-loop regions (**d2**), and cpGFP moiety (**d3**). **d1–d3**, schematics of mutagenesis (top), basal fluorescence images and heatmap images depicting the responses to 100 nM OT (middle), and scatter plots describing the relationship between basal brightness and the fluorescence response to 100 nM OT (bottom). **e**, Traces of fluorescence responses (gray: each trace, green: average trace) of either OT-3.0- or OT-3.0-mut-expressing cells upon stimulation with the indicated drugs. Summary of peak Δ*F*/*F*_0_ (*n* = 10 cells per group). Statistics: one-way ANOVA (*F*_2,27_ = 3.35, *P* = 5.7×10^−16^) with Bonferroni *post-hoc* test (*P* = 4.3×10^−10^: top vs. middle, *P* = 4.3×10^−10^: top vs. bottom, *P* = 1: middle vs. bottom). **f**, Dose–response curves of the sensors (*n* = 10 cells per point). The *F*_max_ and EC_50_ values are summarized on the right. Scale bars, 10 μm (**b, d**). Graphs represent the mean ± SEM (**c, e, f**). \*\*\**P* < 0.001, NS: not significant (**c, e**).

To engineer a fluorescent OT sensor from meOTR, we determined the optimal insertion site of cpGFP in the IL3 of meOTR (Extended Data Fig. 1b). Through the examination of 21 potential mutants expressed in HEK293T cells, a fusion protein in which cpGFP was inserted into the region between K240 and I268 of meOTR (K240−I268) showed the best performance. This fusion protein had robust fluorescence, good PM targeting, and a slight fluorescence response (Δ*F*/*F*_0_ ~ 0.3) to 100 nM OT (Extended Data Fig. 1c–g). Thus, we selected this chimera protein as the template for further engineering.

To improve the fluorescence response of the initial OT sensor, we screened the mutant sensors expressed in HEK293T cells using the following three steps (Fig. 1d). First, we optimized the linkers in the N- and C-terminal regions surrounding the cpGFP. By examining the fluorescence responses of 124 mutants to 100 nM OT stimulation, we obtained OT-1.0, which had an approximately 150% Δ*F*/*F*_0_ response (Fig. 1d1). Next, we extended the mutagenesis to the neighboring regions, ranging from the transmembrane helix to the intracellular loops (TM-to-loop). Based on structural information about the conformational transition during GPCR activation^32,33^, we introduced mutations at sites from position 5.62 in the fifth transmembrane helix (S230 in meOTR) to position 6.36 in the sixth transmembrane helix (M278 in meOTR) (Extended Data Fig. 2a–c). We examined 55 variants, of which five (S230C, F231Y, Q236R, L240G, and T274R in meOTR) showed increased fluorescence responses upon 100 nM OT stimulation. We then developed an improved sensor, OT-2.0, which contained all five of these mutations. OT-2.0 had about a three-fold brighter basal fluorescence and larger fluorescence response (~390% Δ*F*/*F*_0_; Fig. 1d2). Finally, we further developed OT-2.0 by introducing a mutation within the cpGFP. Based on previous knowledge of cpGFP mutagenesis^29^, we screened 34 variants, of which three (S57T, S96M, and N202H in cpGFP; Extended Data Fig. 2b–c) had an increased rate of fluorescence change. We then constructed OT-3.0, a variant that contained all three of these mutations (Fig. 1d3). OT-3.0 had an approximately 720% Δ*F*/*F*_0_ fluorescence response upon stimulation with 100 nM OT, which was suppressed by the OTR antagonist L-368,899 (L-36) (Fig. 1e; top and middle). We also generated an OT-insensitive sensor (OT-3.0 mut) that contained the Y206A mutation, which abolished its ligand-binding capacity^32^ (Fig. 1e; bottom). We characterized the dose-dependent fluorescence responses of OT-1.0−3.0 in HEK293T cells (Fig. 1f), and the results suggested that our screening had successfully improved the dynamic range of fluorescence responses (*F*_max_) with little change to the half maximal effective concentration (EC_50_) values. Herein, we designated the fluorescent module of OT-3.0 as MTRIA (Extended Data Fig. 2b–c) and renamed OT-3.0 as MTRIA_OT_.

We then further characterized the ligand specificity of MTRIA_OT_ expressed in HEK293T cells. Compared with its sensitivity to OT, the sensor was similarly sensitive to isotocin (an OT analog in fish), was much less sensitive to vasopressin orthologs (vasopressin and vasotocin) and inotocin (an OT/vasopressin ortholog in insects), and was insensitive to nematocin (an OT/vasopressin ortholog in nematodes) (Extended Data Fig. 3a–b). We also examined the basic properties of MTRIA_OT_, such as its coupling capacity with downstream effectors, long term-fluorescence stability, and kinetics. We used a Ca^2+^ imaging experiment and a split-luciferase complementation assay to assess the coupling of MTRIA_OT_ with G_αq_ protein and β-arrestin signaling pathways, which are the downstream effectors of the wild type-meOTR. MTRIA_OT_ had no detectable effects on either of these effectors, whereas the wild type-meOTR was able to couple with them (Extended Data Fig. 3c–d). Furthermore, internalization of MTRIA_OT_ from PM was not detected when MTRIA_OT_-expressing HEK293T cells were chronically exposed to 100 nM OT (Extended Data Fig. 3e). Finally, a kinetic analysis of MTRIA_OT_ was conducted using a local puff of OT followed by L-36, which yielded on- and off-rates of approximately 12 s and 5 min, respectively (Extended Data Fig. 3f). Taken together, these results suggest that our MTRIA_OT_ sensor is equipped with enough sensitivity, specificity, and stability to accurately detect extracellular OT dynamics.

### Detection of an OT increase in the brain after exogenous OT administration

Having validated the basic properties of MTRIA_OT_ in HEK293T cells, we next tested whether our OT sensor was applicable to experiments in the brains of living mice. Using an adeno-associated virus (AAV), we expressed MTRIA_OT_ in the anterior olfactory nucleus (AON), a major target site of OT in the brain with a very high OTR expression level^6,7^. We then conducted fiber photometry recordings through an implanted cannula that was placed above the injection site (Fig. 2a and Extended Data Fig. 4a–b). First, we examined the responses of MTRIA_OT_ after the intracerebroventricular infusion of OT in anesthetized mice. We prepared a range of solutions that contained different amounts of OT (0, 0.002, 0.02, 0.2, 2, or 20 μg). When these solutions were serially applied through a stainless cannula implanted into the lateral ventricle (Fig. 2a), significant fluorescence increases were observed upon stimulation with OT at doses of 0.2 μg and above (Fig. 2b–c). This result was consistent with previous findings, that intracerebroventricular infusion of a comparative dose of OT is required to trigger OT-dependent animal behaviors^34–36^. This finding indicates that MTRIA_OT_ is capable of real-time detection of extracellular OT in living brains.

**Figure 2.**
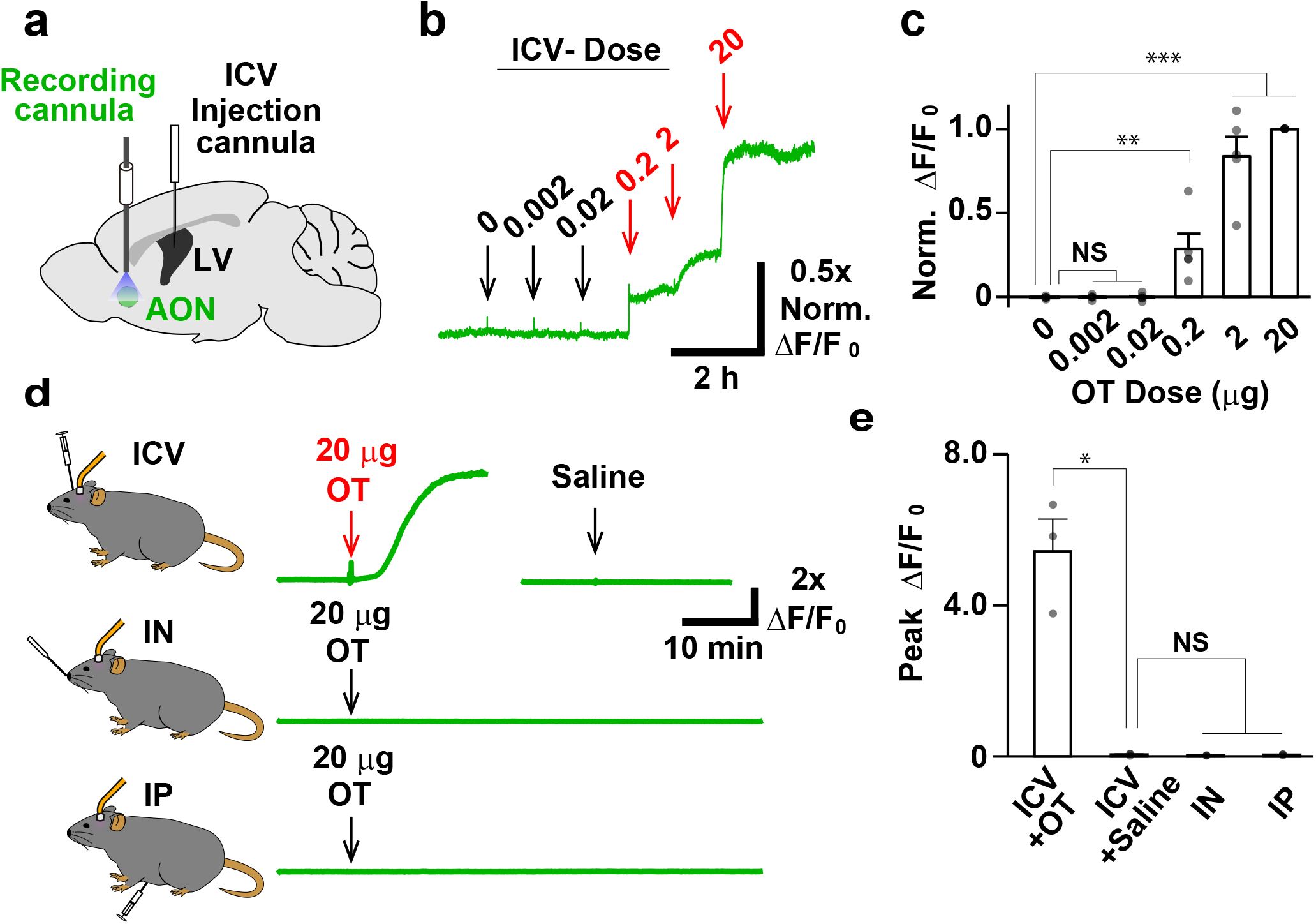
*In vivo* real-time measurement of brain OT dynamics following exogenous OT administration. **a**, Schematic illustrating the fiber photometry recording of MTRIA_OT_ in the AON. **b**, Representative trace of the normalized fluorescence intensity of MTRIA_OT_ following intracerebroventricular (ICV) injection of OT at the indicated dose. **c**, Summary of normalized Δ*F*/*F*_0_. (*n* = 5 mice). Statistics: one-way ANOVA (*F*_5,24_ = 2.62, *P* = 1.4×10^−12^) with Dunnett’s *post-hoc* test to compare with 0 (*P* = 1: 0.002 vs. 0, *P* = 1: 0.02 vs. 0, *P* = 0.0090: 0.2 vs. 0, *P* = 5.4×10^−10^: 2 vs. 0, *P* = 5.2×10^−14^: 20 vs. 0). **d**, Representative traces of Δ*F*/*F*_0_ upon stimulation with either 20 μg OT or saline via the indicated administration routes (top: ICV, middle: intranasal [IN], bottom: intraperitoneal [IP]). **e**, Summary of peak Δ*F*/*F*_0_. (*n* = 3 mice). Statistics: one-way ANOVA (*F*_3,8_ = 4.07, *P* = 3.8×10^−5^) with Bonferroni *post-hoc* test (*P* = 0.020: ICV+OT vs. ICV+saline, *P* = 1: IN vs. ICV+saline, *P* = 1: IP vs. ICV+saline). Graphs represent the mean ± SEM (**c**, **e**). \*\*\**P* < 0.001, \*\**P* < 0.01, \**P* < 0.05, NS: not significant (**c**, **e**).

In the past decade, whether the peripheral administration of OT can alter brain OT levels has been controversial^17^. Because it has been reported that the peripheral administration of less than 20 μg of OT is sufficient to affect animal behaviors^37,38^, we next evaluated whether OT levels in the brain increased after the application of exogenous OT from two distinct peripheral administration routes: intranasal or intraperitoneal. Neither intranasal nor intraperitoneal administration of high concentrations of OT induced significant fluorescence responses of MTRIA_OT_ in anesthetized mice (Fig. 2d–e). Taken together with the finding that MTRIA_OT_ even had robust responses to the intracerebroventricular administration of OT solution diluted at 1:100 (Fig. 2b–c), out data suggest that the blood–brain barrier permeability to OT is less than 1%.

### OT oscillation in the brains of freely behaving mice

Having shown that MTRIA_OT_ is functional in the mouse brain, we next examined whether MTRIA_OT_ can be used to assess endogenous OT dynamics in the brains of adult mice. We virally expressed MTRIA_OT_ in the AON and measured the fluorescence responses using a fiber photometry technique (Fig. 3a). Surprisingly, MTRIA_OT_-mediated fluorescence measurements revealed that transient OT signals were repeated at approximately 2-hour intervals when mice were freely behaving in a cage with food and water supplied *ad libitum* (Fig. 3b–d). We named this ultradian OT rhythm “OT oscillation.” OT oscillations were absent in simultaneously recorded reference signals (405 nm of MTRIA_OT_; Fig. 3b; left bottom) and in signals recorded using control sensors (470 nm and 405 nm of MTRIA_OT_-mut; Fig. 3b; right), excluding the involvement of artifacts derived from movements and/or autofluorescence. The expression of tetanus toxin light chain, which prevents vesicular transmitter release^39^, in OT-expressing neurons in the PVN suppressed the increase in OT compared with the control (Fig. 3e–h), confirming that OT oscillation is dependent on OT release from oxytocinergic neurons in PVN.

**Figure 3.**
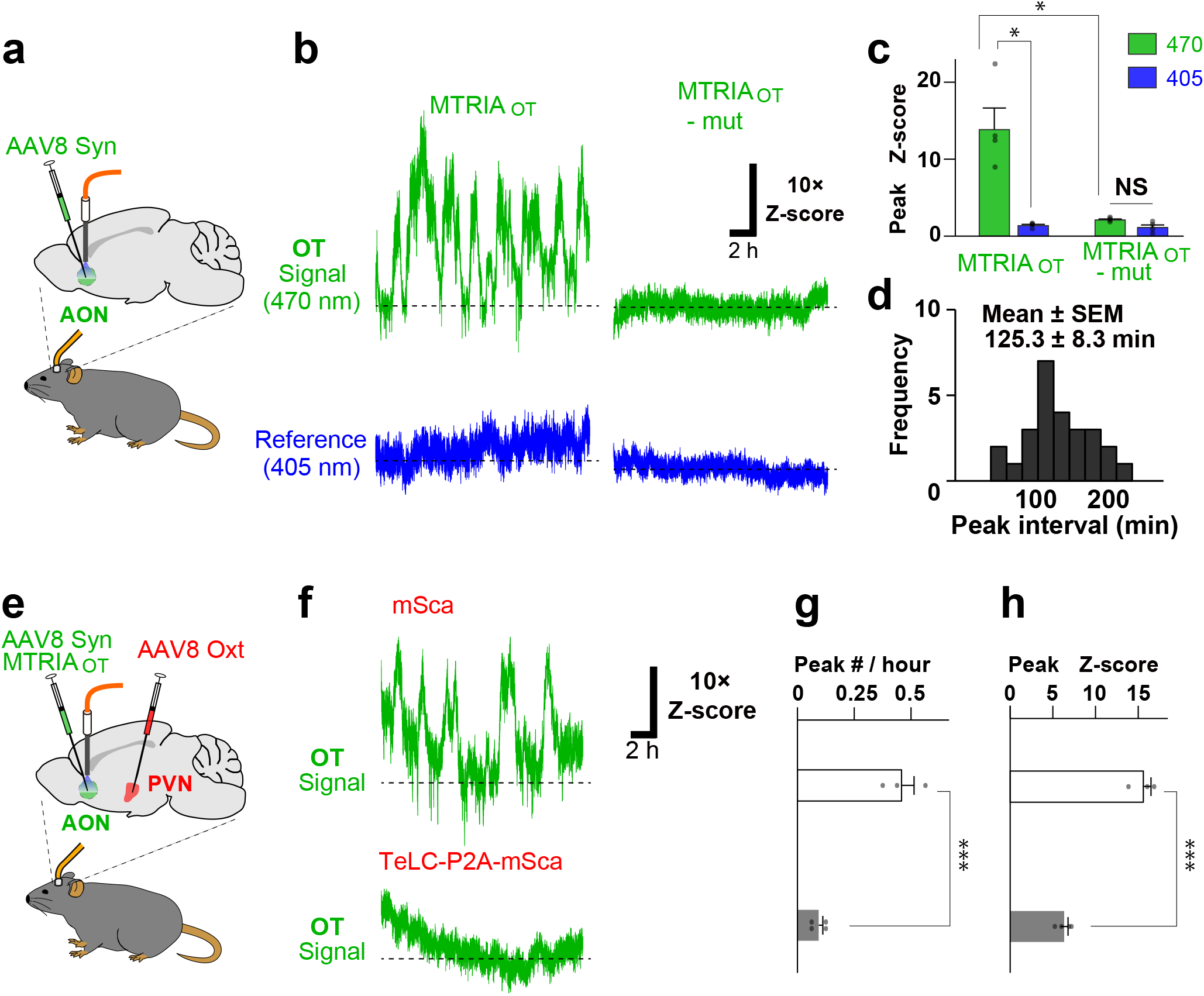
*In vivo* real-time measurement of brain OT dynamics in freely behaving mice. **a**, Schematic illustrating the fiber photometry recording of MTRIA_OT_ in the AON in freely behaving mice. **b**, Representative traces of 470 nm-excited signals (green) and 405 nm-excited signals (blue) from either MTRIA_OT_- or MTRIA_OT_ mut-expressing mice. **c**, Summary of peak z-scores (*n* = 4 mice). Statistics: one-way ANOVA (*F*_3,12_ = 3.49, *P* = 7.4×10^−5^) with Bonferroni *post-hoc* test (*P* = 0.026: 470 nm vs. 405 nm in MTRIA_OT_, *P* = 0.034: 470 nm in MTRIA_OT_ vs. 470 nm in MTRIA_OT_-mut, *P* = 0.21: 470 nm vs. 405 nm in MTRIA_OT_-mut). **d**, Frequency histogram showing the intervals of OT signal peaks (*n* = 26 events from four mice). **e**, Schematic illustrating the experimental protocol for assessing the involvement of vesicular release from PVN OT neurons in the OT signal increase in the AON. **f**, Representative traces of MTRIA_OT_ activities recorded from mice either expressing mSca or co-expressing tetanus toxin light chain (TeLC) and mSca in the PVN. **g**, **h**, Summary of the peak number every hour (**g**) and peak z-score (**h**) (*n* = 3 mice in mSca and *n* = 4 mice in TeLC-P2A-mSca). Statistics: unpaired two-tailed *t*-test (*P* = 8.2×10^−4^ in **g** and *P* = 1.6×10^−4^ in **h**). Graphs represent the mean ± SEM (**c**, **g**, **h**). \*\*\**P* < 0.001, \**P* < 0.05, NS: not significant (**c**, **g**, **h**).

We next explored what might affect the patterns of OT oscillation. Because the sleep–wake cycle is a key regulator of biological rhythm, we evaluated OT oscillation in mice when sleep was induced by anesthesia. When an anesthetic mixture of dexmedetomidine, butorphanol, and midazolam was intraperitoneally administered, the fluorescence signal of MTRIA_OT_ fell to a level below the baseline (Fig. 4a, b), suggesting that brain OT levels are suppressed by anesthesia.

**Figure 4.**
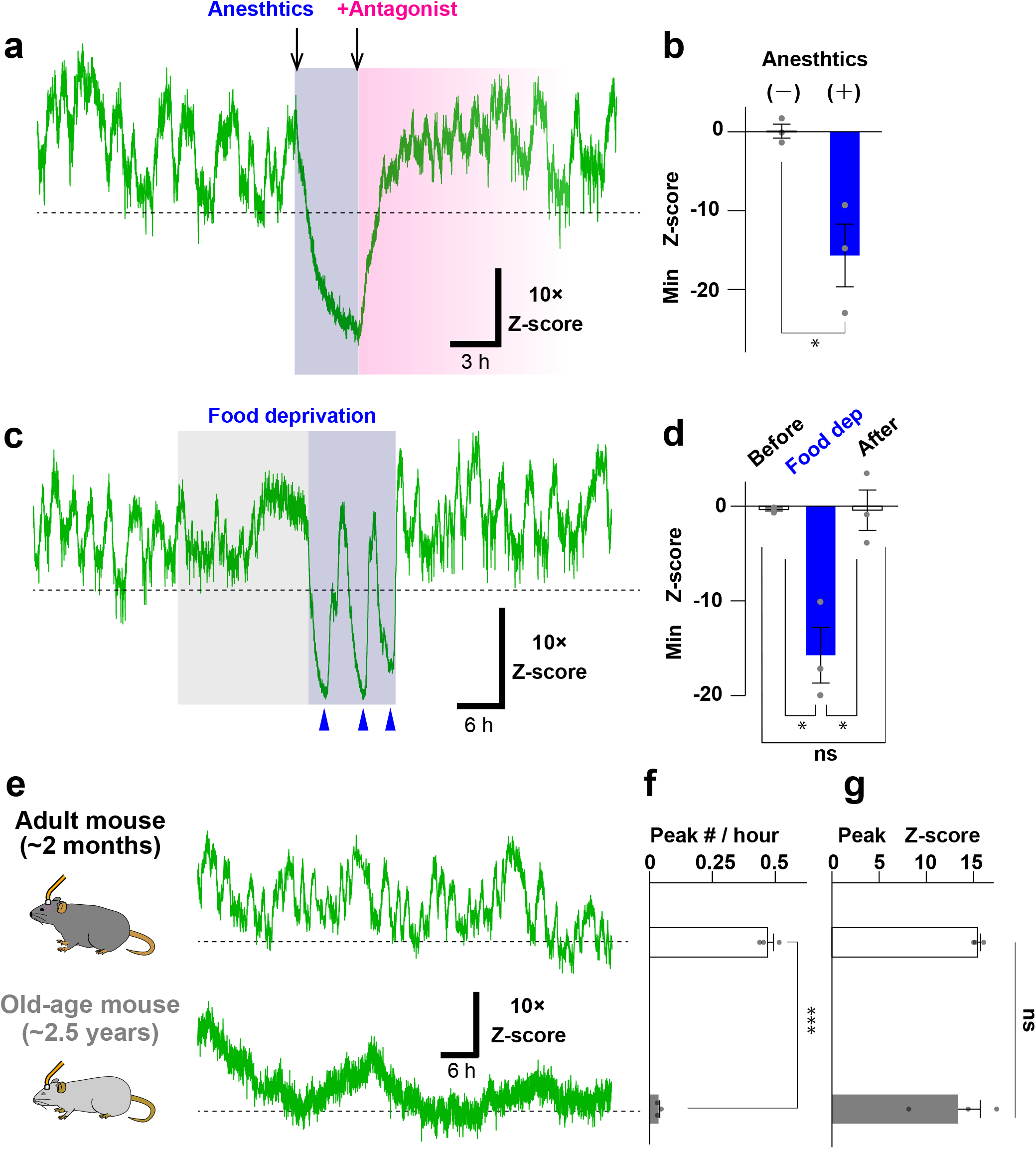
Alterations in OT oscillation caused by anesthesia, food-deprivation, and aging. **a**, Representative trace of MTRIA_OT_ activity showing the impact of anesthesia on OT oscillation. The background of the trace is shaded to indicate the period during anesthesia (dark blue) and the period after release by an antagonist (pink). **b**, Summary of the minimum values of z-scores before and during anesthesia (*n* = 3 mice). Statistics: unpaired two-tailed *t*-test (*P* = 0.018). **c**, Representative trace of MTRIA_OT_ activity showing the impact of food deprivation on OT oscillation. The background of the trace is shaded to indicate the period of food deprivation. The period of OT turbulence is colored dark blue and the peaks of undershot signal are indicated by arrowheads. **d**, Summary of the minimum z-score values before, during (Food dep.), and after food deprivation (*n* = 3 mice). Statistics: one-way ANOVA (*F*_2,6_ = 5.14, *P* = 0.0030) with Bonferroni *post-hoc* test (*P* = 0.019: before vs. Food dep., *P* = 0.040: Food dep. vs. after, *P* = 1: before vs. after). **e**, Schematic illustrating the recordings and representative traces of MTRIA_OT_ fluorescence signals in adult (top) and old-aged (bottom) mice. **f**, **g**, Summary of the peak number every hour (**f**) and peak z-score (**g**) (*n* = 3 mice). Statistics: unpaired two-tailed *t*-test (*P* = 6.0×10^−5^ in **f** and *P* = 0.49 in **g**). Graphs represent the mean ± SEM (**b**, **d**, **f**, **g**). \*\*\**P* < 0.001, \**P* < 0.05, NS: not significant (**c**, **g**, **h**).

Given that the feeding process is reportedly associated with OT release^40^, we examined the impact of fasting stress on brain OT dynamics. Interestingly, after about half a day of food deprivation, the patterns of OT oscillation gradually became disturbed; the oscillatory signals undershot to a level below the initial baseline (Fig. 4c, d). We named this phenomenon “OT turbulence” (Fig. 4c; dark blue-shaded region). After refeeding, this OT turbulence halted and normal OT oscillation quickly recovered (Fig. 4c, d). This result suggests that fasting stress is likely to be encoded as OT turbulence in the brain.

Because aging is associated with a decline in the neuroendocrine system^15,41^, we next analyzed the differences in AON OT dynamics between adult (~2 months old) and old-aged (~2.5 years old) mice. The fiber photometry-mediated fluorescence measurements revealed the occurrence of oscillatory OT responses in both groups, but the frequency of OT transients became much slower in the old-aged animals (~24-hour intervals) compared with those in adult animals (~2-hour intervals), although there were no significant changes in amplitudes (Fig. 4e, f). These results suggest that an altered frequency of OT oscillation may underlie aging-associated declines in brain function.

### Development of fluorescent sensors for various ligands by conjugating MTRIA to other GPCRs

Because MTRIA was able to successfully produce a highly sensitive optical readout of OT-dependent OTRs, we finally examined whether MTRIA was also able to detect ligand binding-induced conformational changes of other GPCRs (Fig. 5a). We cloned 184 receptors for 46 ligands derived from either the human, mouse, zebrafish, or medaka, and conjugated MTRIA to a region ranging from position 5.62 of TM5 to position 6.36 of TM6 in the receptors (Extended Data Fig. 2a). To our surprise, almost 30% of the engineered sensors (54/184 proteins) showed a marked fluorescence increase (> 50% Δ*F*/*F*_0_) upon stimulation with high concentrations of their specific ligand (Fig. 5b). We listed the 24 sensors that showed the largest fluorescence response among the sensors sharing the same ligand (Fig. 5c, Extended Data Fig. 5) and named them MTRIA sensors (e.g., a sensor for dopamine was called MTRIA_DA_). These results demonstrate the utility of MTRIA for the efficient development of new GPCR-based fluorescent sensors. This simple and efficient approach, MTRIA system, will contribute to accelerating the engineering of various GPCR-based sensors.

**Figure 5.**
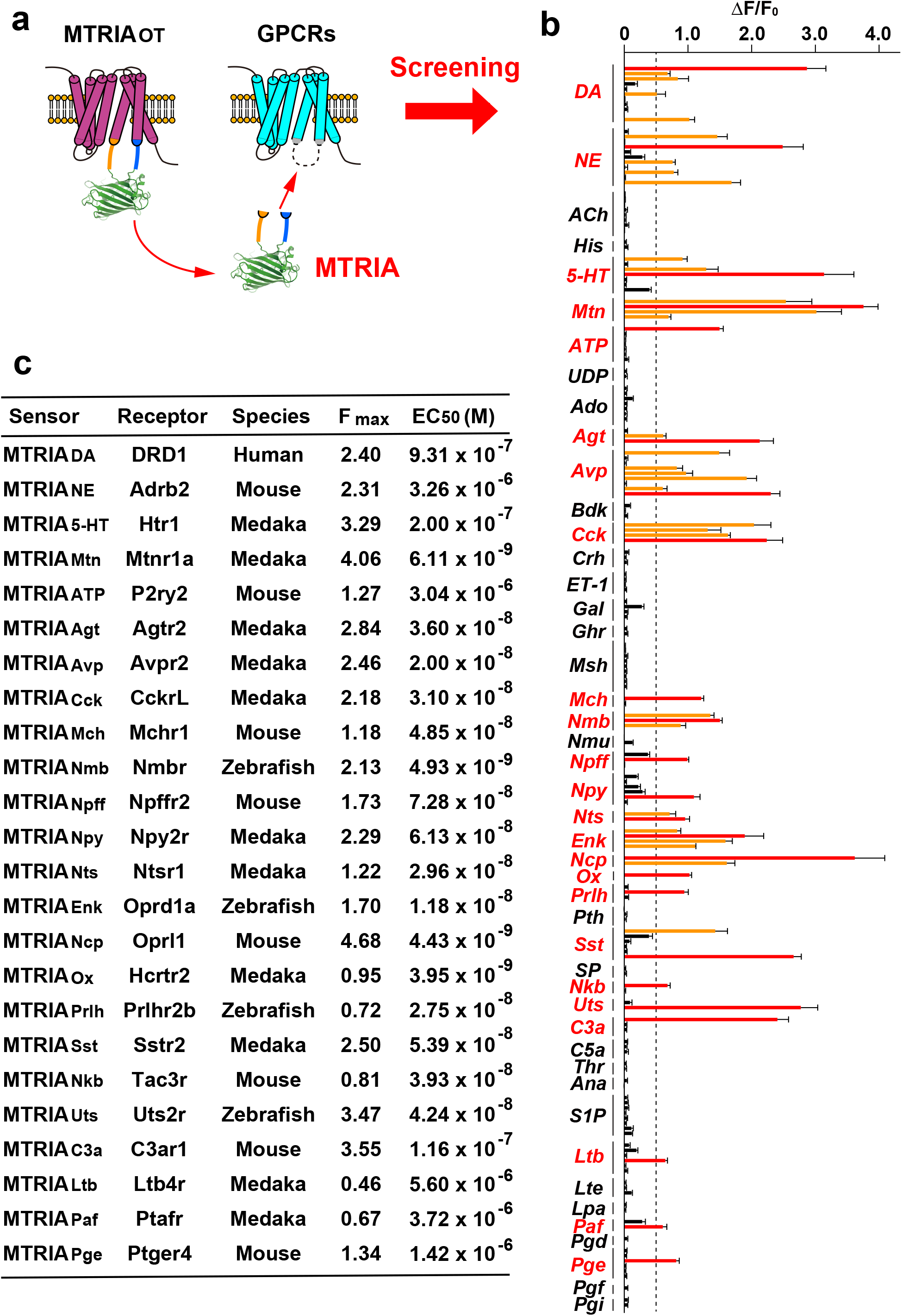
Development of various transmitter sensors by conjugation with MTRIA. **a**, Schematic illustrating the development of MTRIA sensors. **b**, The Δ*F*/*F*_0_ values are given as the mean ± SEM (*n* = 5 cells for each group). The sensors with Δ*F*/*F*_0_ > 0.5 are colored either red or orange. The sensors with the best performance among receptors sharing the same ligand are colored red. **c**, Table showing the basic properties of the MTRIA sensors: sensor names, scaffold receptors, species, *F*_max_, and EC_50_.

## DISCUSSION

In this study, we described the development and application of a new fluorescent sensor for OT, MTRIA_OT_, which displayed excellent properties for measuring extracellular OT; it had a large dynamic range (~720% Δ*F*/*F*_0_), an optimal affinity to OT (EC_50_ of ~20 nM), ligand specificity to OT orthologs, minimal effects on cellular signaling, and long-term fluorescence stability. We then demonstrated the applications of MTRIA_OT_ for measuring exogenously applied and endogenous OT dynamics in the living brain. We also described the utility of the fluorescent module of MTRIA_OT_ (i.e., MTRIA) for the efficient development of functional GPCR-based fluorescent sensors for various ligands. Our new sensor will be valuable for analyzing the dynamics of extracellular signaling *in vivo*.

Although the impacts of exogenous OT administration on mood and emotions are expected to lead to new treatments for psychiatric disorders, direct evidence for an increase in brain OT levels following peripheral OT administration remains limited. Here, we demonstrated that MTRIA_OT_-mediated OT recording is useful for detecting changes in brain OT levels following exogenous OT administration. In the present study, neither intranasal nor intraperitoneal administration of a relatively large amount (20 μg) of OT resulted in significant changes in the fluorescence signal of MTRIA_OT_ in the brain of adult mice. In contrast, intracerebroventricular administration of OT at concentrations as low as 0.2 μg triggered a robust increase in brain OT levels (Fig. 2), supporting the notion that the efficacy of OT transport through the blood–brain barrier is very low^42^. As has been mentioned elsewhere^17,43^, invasive measurements of OT using acutely implanted probes may lead to the leakage of OT from injured blood vessels into the brain parenchyma following peripheral OT administration. Given that peripheral OTRs are distributed in various tissues including the heart, adrenal medulla, adipose tissue, and peripheral neurons^3^, the actions of exogenously applied OT on peripheral OTRs may have an impact on animal behaviors.

Accumulating evidence suggests that rhythmic hormonal signaling, with frequencies ranging from a few hours (ultradian) or around a day (circadian) to more than a day (infradian), plays an important role in the regulation of a range of biological processes, including body homeostasis (e.g., sleep, feeding, and metabolism), mood, and cognitive performance^44–46^. Although circadian changes in OT levels in the cerebrospinal fluids of primates^12,47^ and rodents^48^ have been proposed, it has not yet been explored whether ultradian regulation of OT levels exist; this is presumably because of the limited sensitivity and slow sampling rates of previous techniques. Taking advantage of the faster time resolution of our time-lapse fluorescence recordings, we revealed that OT levels in the AON exhibit ultradian rhythms with approximately 2-hour intervals in freely behaving mice. However, we cannot exclude the existence of much faster OT transients, which may be masked by the slow kinetics of MTRIA_OT_ in fluorescence decay (*τ*_off_ > 10 min) upon agonist dissociation. Our findings lead to a question: how is OT oscillation regulated? It should be noted that PVN neurons in slice cultures show spontaneous synchronized oscillations in Ca^2+^ activity at intervals of a few hours^49^, implicating that ultradian OT rhythms may be controlled by a Ca^2+^-dependent biological clock. There may also be rhythmic changes in other neuromodulators that show synchronization with OT oscillation. It is of particular interest that ultradian rhythms of glucocorticoid secretion, which is thought to be coupled with OT levels^50^, is important for good sleep quality and cognitive function in healthy humans^44^. Thus, future efforts are needed to clarify the regulatory mechanisms and physiological significance of OT oscillation.

Overall, our MTRIA_OT_-mediated measurements suggest the importance of temporal information about OT dynamics in the brain. Moreover, our analyses indicate that the patterns of ultradian OT oscillation can be affected by anesthesia, food deprivation, and aging. Thus, it is important to carefully consider the experimental conditions and the state of subjects when interpreting OT measurement data in the brain. The context- and subject-dependent inconsistencies regarding the effects of OT in human clinical trials^20^ may be related to such parameters. The use of MTRIA_OT_ will allow us to extend our knowledge of brain OT dynamics and their associations with complex behaviors.

## METHODS

### DNA constructions

DNA encoding GPCRs, a red fluorescent Ca^2+^ sensor jRGECO1a^51^, and PM-targeted mScarlet^52^ (mSca_mem_) were amplified by polymerase chain reaction (PCR), and were then subcloned into the pCIS vector, which is a CAG promoter-driven expression vector^53^. The template DNA for GPCRs was prepared using the following four approaches. First, cDNA libraries of a mouse (*Mus musculus*) brain, zebrafish (*Danio rerio*) brain, and medaka (*Oryzias latipes*) brain were prepared using the RNeasy Kit (Qiagen, Hilden, Germany) and QuantiTect Reverse Transcription Kit (Qiagen). Second, genomic DNAs of human (*Homo sapiens*) and mouse were isolated from HEK293T cells and a mouse brain, respectively. Third, gene fragments of OTRs derived from chicken (*Gallus gallus*), snake (*Notechis scutatus*), and frog (*Xenopus laevis*) were synthesized by Integrated DNA Technologies (Coralville, IA, USA). Finally, for cloning of human dopamine receptor DRD1, human muscarinic receptor CHRM3, and human adrenergic receptor ADRA2A, plasmids encoding dLight1.1, GACh2.0, and GRAB_NE_1m were used as the templates, respectively^27–29^. GPCRs used in Fig. 5 and the primer pairs for their cloning were listed in Extended Data Table 1. To assess the localization of OTRs in HEK293T cells, a human influenza hemagglutinin (HA)-tag was added to the N-terminal regions of the OTRs using PCR. For the construction of the green-fluorescent transmitter sensors, cpGFPs derived from GCaMP6s were inserted into the IL3 regions of GPCRs using either overlap extension PCR^54^ or the Gibson assembly protocol^55^. For mutant sensor screening, site-directed mutagenesis was performed using primers containing mutated nucleotide(s). In particular, randomized site-directed mutagenesis was achieved using primers containing randomized codons (NNN). For the `construction` of plasmid DNA in the β-arrestin recruitment assay, fragments of NanoLuc—SmBit and LgBit— were amplified from pNL1.1 (Promega, Madison, WI, USA). They were then fused with the C-terminus of OTR and the N-terminus of rat β-arrestin^56^, followed by cloning into the pCIS vector. For the construction of transfer plasmid DNA for AAVs, MTRIA_OT_ and MTRIA_OT_-mut were cloned into the pAAV Syn woodchuck hepatitis virus post-transcriptional regulatory element (WPRE) vector, which is a plasmid driven by a human synapsin promoter. In addition, mScarlet (mSca) and tetanus toxin light chain-P2A-mSca were cloned into the pAAV Oxt WPRE vector, which is a plasmid driven by a mouse OT promoter.

### Cell culture

HEK293T cells were cultured in Dulbecco’s Modified Eagle’s Medium (FUJIFILM-Wako, Osaka, Japan) supplemented with 10% fetal bovine serum (Thermo Fisher Scientific, Waltham, MA, USA), 1007U/7mL penicillin, and 1007μg/mL streptomycin at 37°C under a humidified atmosphere of 5% CO_2_ and 95% air. For the fluorescence microscopy observations, cells were transfected with 1 μg of plasmid using 2 μg of Polyethylenimine Max (PEI MAX) transfection reagent (Polysciences, Warrington, PA, USA), and were seeded onto either glass bottom dishes or glass bottom chambers (Matsunami, Osaka, Japan) coated with collagen (Nitta Gelatin, Osaka, Japan). After a 3-hour incubation, the medium was changed, and the cells were further cultured for 24–36 hours before the imaging experiments.

### AAV production

pHelper, XR8, and a transfer plasmid were simultaneously transfected into HEK293T cells using PEI MAX transfection reagent. After overnight incubation, the medium was changed. Supernatant was collected at 48- and 96-hour time points, after medium exchange. Viral particles were purified using a polyethylene glycol-mediated precipitation method^57^. The purified viral particles were then concentrated using an ultrafiltration membrane unit (Amicon Ultra; EMD Millipore, Burlington, MA, USA). Virus titers were determined by quantitative PCR using a pair of primers for the WPRE sequence (forward: actgtgtttgctgacgcaac, reverse: agcgaaagtcccggaaag). Viral vectors with titers > 10^12^ gc/mL were used for the fiber photometry measurements.

### Immunostaining of transfected cells

For the immunostaining analysis of the cellular localization of OTRs, HEK293T cells that had been co-transfected with an HA-tagged OTR and mSca_mem_ (10:1 ratio) were fixed in 4% (w/v) paraformaldehyde-containing phosphate-buffered saline (PBS; FUJIFILM-Wako) for 10 min at room temperature. Next, cells were permeabilized and blocked for 30 min at room temperature in blocking solution (PBS containing 0.2% TritonX-100 [PBST] and 5% normal goat serum [FUJIFILM Wako]), and were then incubated with anti-HA antibody (rabbit; MBL, Aichi, Japan) for 30 min at room temperature. After washing with PBST, samples were incubated with Alexa 488-conjugated anti-rabbit IgG antibody (goat; Jackson ImmunoResearch, West Grove, PA, USA) for 30 min at room temperature. After three washes with PBST, the stained cells in PBS underwent microscopic analysis.

### Sensor screening and evaluation by fluorescence microscopy

The fluorescence images in Figures 1 and 5 and Extended Data Figures 1, 3, 4, and 5 were obtained using Dragonfly 301, a spinning-disk confocal microscope system (Andor Technology, Belfast, UK) equipped with a Nikon Eclipse Ti2 (Nikon, Tokyo, Japan), lasers (405, 488, 561, and 637 nm), emission filters (450 ± 25 nm for the 405 nm laser, 525 ± 25 nm for the 488 nm laser, 600 ± 25 nm for the 561 nm laser, and 700 ± 37.5 nm for the 637 nm laser), and an electron multiplication charge-coupled device (EMCCD) camera (iXon Life 888; Andor Technology). Either a dry objective lens (CFI PLAN APO 20×, NA 0.75; Nikon) or a water-immersion lens (CFI Plan Apo IR 60×C WI, NA 1.27; Nikon) was used to illuminate the lasers. For the live-imaging experiments, data were acquired every 5 s. To analyze the cellular localization of fluorescent sensors, z-stack images were acquired at 0.5 μm steps. For the screening of fluorescent sensors, transfected cells seeded onto glass-bottomed chamber slides were soaked in 200 μL artificial extracellular solution (ECS; 150 mM NaCl, 4 mM KCl, 2 mM CaCl_2_, 1 mM MgCl_2_, 5 mM HEPES, and 5.6 mM glucose; pH 7.4). They were then stimulated with 200 μL drug-containing ECS, in which the drug concentration was twice the final concentration, administered through a pipette. To analyze the ligand dose–response of the fluorescent sensors, time-constant measurements, Ca^2+^ imaging, and the evaluation of long-term signal stability were performed. The transfected cells were seeded onto a collagen-coated glass-bottomed dish and continuously perfused with ECS. They were then rapidly stimulated with drug-containing solution using a custom-made perfusion system equipped with solenoid valves.

### β-Arrestin recruitment assay

The recruitment of β-arrestin to meOTR and MTRIA_OT_ was assessed using the split-luciferase complementation assay. Either meOTR-SmBit or MTRIA_OT_-SmBit was co-transfected with LgBit-arrestin in HEK293T cells that were seeded onto an opaque 96-well plastic plate using PEI MAX reagent. Twenty-four hours after transfection, the cells were washed with Hanks’ balanced salt solution (HBSS), and stimulated with HBSS containing 100 nM OT. After the addition of a luciferase substrate (ONE-Glo™ Luciferase Assay System, Promega), luminescence was measured using a plate reader (Infinite 200 Pro F Plex; Tecan, Männedorf, Switzerland).

### Reagents

OT, angiotensin (Agt), [Arg8]-vasopressin (Avp), bradykinin (Bdk), cholecystokinin (Cck), corticotropin-releasing hormone (Crh), endothelin-1 (ET-1), galanin (Gal), ghrelin (Ghr), α-melanocyte stimulating hormone (Msh), melanin-concentrating hormone (Mch), neuromedin B (Nmb), neuromedin U (NmU), neuropeptide Y (Npy), neurotensin (Nts), met-enkephalin (Enk), nociceptin (Ncp), orexin B (Ox), prolactin-releasing hormone (Prlh), parathormone (Pth), somatostatin (Sst), substance P (SP), neurokinin B (Nkb), and urotensin (Uts) were purchased from Peptide Institute (Osaka, Japan). L-368,899 hydrochloride, neuropeptide ff (Npff), sphingosine-1-phosphate (S1P), uridine 5′-diphosphate disodium salt (UDP), and anandamide (Ana) were purchased from Tocris Bioscience (Bristol, United Kingdom). Isotocin, C3a anaphylatoxin (C3a), and C5a anaphylatoxin (C5a) were purchased from Bachem (Bubendorf, Switzerland). Vasotocin and thrombin receptor agonist (Thr) were purchased from Anygen (Gwangju, Korea), and inotocin was purchased from Phoenix Pharmaceuticals (Mannheim, Germany). Histamine disodium salt (His) and melatonin (Mtn) were purchased from FUJIFILM-Wako. Leukotriene B4 (Ltb), leukotriene E4 (Lte), 1-oleoyl lysophosphatidic acid (LPA), platelet-activating factor C-16 (PAF), and prostagrandin F2a (Pgf) were purchased from Cayman Chemicals (Ann Arbor, MI, USA). Prostagrandin D2 (Pgd), prostagrandin E2 (Pge), prostagrandin I2 sodium salt (Pgi), 5-hydroxytryptamine hydrochloride (5-HT), dopamine hydrochloride (DA), acetylcholine chloride (ACh), and adenosine 5′-triphosphate disodium salt (ATP) were purchased from Nacalai (Kyoto, Japan). Epinephrine (Epi) was purchased from Daiichi-Sankyo (Tokyo, Japan), and norepinephrine (NE) was purchased from Alfresa Pharma Corporation (Osaka, Japan). Nematocin was synthesized by Cosmo Bio (Tokyo, Japan). The final concentration of the applied ligands in Fig. 5b were as follows;

100 μM: DA, NE, ACh, His, 5-HT, ATP, UDP, and Ado; 1 μM: Enk, C3a, Ana, Lpa, Paf, Pgd, Pge, Pgf, and Pgi; and 100 nM: Mtn, Agt, Avp, Bdk, Cck, Crh, ET-1, Gal, Ghr, Msh, Mch, Nmb, Nmu, Npff, Npy, Nts, Ncp, Ox, Prlh, Pth, Sst, SP, Nkb, Uts, C5a, Thr, S1P, Ltb, and Lte.

### Animal surgery

All animal procedures were conducted in accordance with the guidelines of Kanazawa University and Osaka University. C57BL/6 J or C57BL/6N female mice at either 6–8 weeks postnatal (adult mice) or 2.5 years postnatal (old-age mice) were purchased from CLEA Japan Inc. (Tokyo, Japan). Stereotaxic surgery was performed under anesthesia with isoflurane. The depth of anesthesia was assessed using the tail pinch method. Body temperature was maintained using a heating pad. One microliter of AAV suspension within a glass micropipette was injected into the left medial AON (2.2 mm anteroposterior [AP] and 0.3 mm mediolateral [ML] from bregma, and −4.0 mm dorsoventral [DV] from the skull surface). After the virus injection, a fiber-optic cannula with a 400 μm core diameter and 0.39 NA (CFMC14L05; Thorlabs, Newton, NJ, USA) was implanted just above the injection site (2.2 mm AP, 0.3 mm ML, and −3.8 mm DV). For the intracerebroventricular injection experiments, a stainless cannula fabricated from a 22 G needle was additionally inserted into the right lateral ventricle (−0.7 mm AP, 1.5 mm ML, and −2.5 mm DV). To fix and protect the implanted cannula(s), dental cement (Ketac Cem Easymix; 3M, Maplewood, MN, USA) and silicone rubber (Body Double; Smooth-On, Macungie, PA, USA) were used. Carprofen (5 mg/kg; intraperitoneal), a non-steroidal anti-inflammatory drug, and buprenorphine (0.1 mg/kg, intraperitoneal), an opioid analgesic, were administered after the surgery. At 2 weeks or more following the AAV injection, the mice were used for the fiber photometry experiments. On the measurement day, the cannula was coupled to a patch cable (M79L01; Thorlabs) via an interconnector (ADAF1 or ADAF2; Thorlabs).

### Fiber photometry measurements

The fiber photometry setup (Extended Data Fig. 4a) was constructed as described previously, with a minor modification^58^. Excitation light from a light-emitting diode (LED) was directed into a patch cable (M79L01) through an objective lens (CFI Plan Fluor 20× lens; Nikon), and the emission light was projected onto the sensor of a scientific complementary metal-oxide semiconductor (sCMOS) camera (Zyla 4.2P; Andor Technology) after passing through a dichroic mirror (Di01-R405/488/561/635-25×36; Semrock, Rochester, NY, USA) and an emission filter (YIF-BA510-550S; SIGMA KOKI, Tokyo, Japan). To acquire an OT-dependent signal and an isosbestic signal, two different LED light sources—a 470 nm light (light from M470F3 filtered with FB470-10, 4 μW at the fiber tip; Thorlabs) and a 405 nm light (light from M405FP1 filtered with FB410-10, 4 μW at the fiber tip; Thorlabs)—were alternatively switched. Data were acquired at 2 s intervals with an LED light exposure of 0.3s. The device control and data acquisition were conducted using a custom-made LabVIEW program (National Instruments, Austin, TX, USA). Data were processed using Fiji software^59^, and were then analyzed using a custom-made Python program. The experiments were conducted within a cage (width: 270 mm, length 440 mm, and height 18.7 mm), and the behaviors of mice were captured every 1 s using an overhead camera (BFS-U3-I3Y3-C; FLIR Systems, Wilsonville, OR, USA). To visualize the behaviors of mice in the dark environment, light from an infrared LED array (AE-LED56V2; Akizuki, Aichi, Japan) was used. Unless stated otherwise, mice were fed standard pellet and water *ad libitum*. For the experiments shown in Fig. 2 and Fig. 4a, mice were anesthetized by the intraperitoneal administration of a mixture of 0.375 mg/kg dexmedetomidine hydrochloride, 2 mg/kg midazolam, and 2.5 mg/kg butorphanol tartrate. For recovery from anesthesia, 0.75 mg/kg atipamezole hydrochloride was applied intraperitoneally.

### Verification of implantation sites

After the fiber photometry measurements, we confirmed the sensor expression and cannula insertion sites by immunostaining. The brains of anesthetized mice were rapidly removed after decapitation, and were immersed in 4% (w/v) paraformaldehyde-containing PBS overnight at 4°C, followed by incubation in 20% (w/v) sucrose-containing PBS overnight at 4°C. After being embedded in optimal cutting temperature compound (Sakura Finetek, Tokyo, Japan), coronal sections of brains were prepared at 50 μm thickness using a cryostat (CM1950; Leica Microsystems, Wetzlar, Germany). For immunostaining, the sections were permeabilized in blocking solution for 30 min at room temperature, and were then incubated with anti-GFP antibody (rabbit; MBL; 1:1000 dilution) overnight at 4°C. After three washes with PBST, the sections were incubated with Alexa 488-conjugated anti-rabbit IgG antibody (goat; Jackson ImmunoResearch; 1:2000 dilution) for 40 min at room temperature. After three washes with PBST and three washes with PBS, the samples were mounted in polyvinyl alcohol-based mounting medium containing 1 μg/mL 4′,6-diamidino-2-phenylindole (DAPI; FUJIFILM-Wako) and 2.5% (w/v) 1,4-diazabicyclo[2.2.2]octane (Sigma, St. Louis, MO, USA) as an antifade. The stained sections were then imaged using a spinning-disk confocal microscope system (Dragonfly 301). Mice with misplacement of the cannula were excluded from the analysis.

### Statistical analysis

All summary data are expressed as the mean ± standard error of the mean (SEM) unless stated otherwise. The z-score was calculated as the average peak response divided by the standard deviation of the baseline fluorescence. For the comparison of two groups, we used two-tailed Student’s *t*-tests. One-way analysis of variance (ANOVA) tests were performed when more than two groups were compared, followed by either the Tukey–Kramer or Dunnett’s *post-hoc* test. Throughout the study, *P* < 0.05 was considered statistically significant. Data distribution was assumed to be normal, and variance was similar between the groups that were statistically compared. No statistical methods were used to predetermine the sample sizes, but our sample sizes were similar to those generally used in the field^26–30^.

## ACKNOWLEDGEMENTS

We thank the members of the Nishiyama and Hibino laboratories for their helpful discussions, Tambo for technical assistance, D. Gadella (University of Amsterdam) for sharing pmScarlet_C1 (Addgene plasmid #85042), D. Kim and the GENIE Project (Janelia Research Campus) for sharing pGP-CMV-NES-jRGECO1a (Addgene plasmid # 61563), Y. Li (Peking University) for sharing pAAV-hSyn-GRAB_NE1m (Addgene plasmid # 123308) and pDisplay-GACh2.0 (Addgene plasmid # 106073), L. Tian (UC Davis) for sharing pCMV-dLight1.1 (Addgene plasmid # 111052), and R. Lefkowitz (Duke University) for sharing β-arrestin2 GFP WT (Addgene plasmid # 35411). This work was supported by grants from the Ministry of Education, Culture, Sports, Science and Technology to D.I. (18K15036); Japan Agency for Medical Research and Development to M.N. (JP19gm6310008); Takeda Science Foundation to D.I.; LOTTE Foundation to D.I.; Research Foundation for Opto-Science and Technology to D.I.; Konica Minolta Science and Technology Foundation to D.I.; Salt Science Research Foundation to D.I.; and Hokuriku Bank to D.I.

## AUTHOR CONTRIBUTIONS

D.I. conceived the project, performed the experiments, and analyzed the data. D.I., H.H., and M.N. discussed the results and wrote the manuscript.

## COMPETING INTERESTS

The authors declare no competing financial interests.

## DATA AVAILABILITY

The data that support the findings of this study are available from the corresponding author upon reasonable request.

## CODE AVAILABILITY

All analysis scripts are available from the corresponding author upon reasonable request.

## EXTENDED DATA FIGURE LEGENDS

**Extend Data Figure 1.**
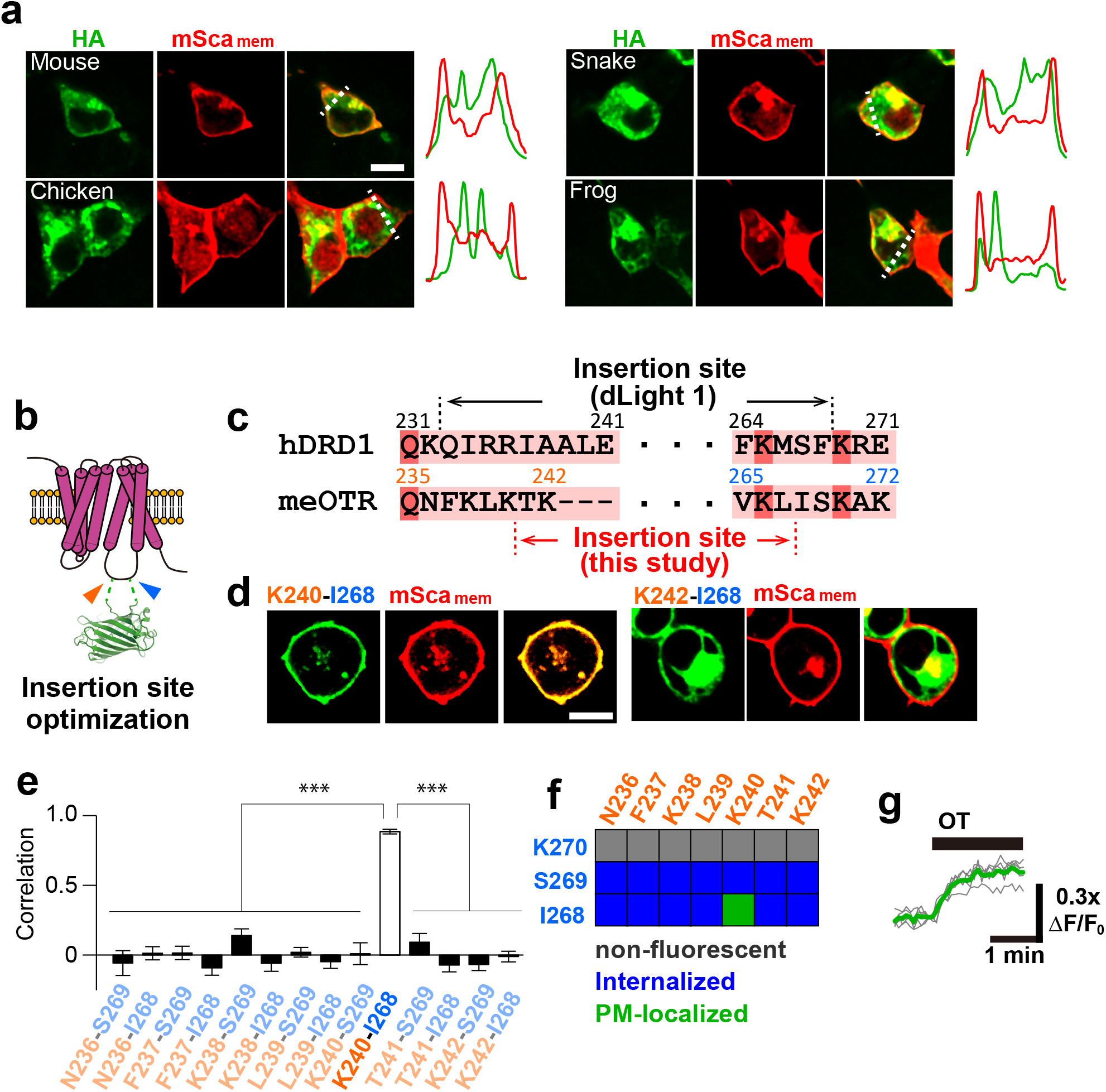
Optimization of a new fluorescent OT sensor. **a**, Representative images of HEK293T cells co-expressing an HA-tagged OTR (green) and a mSca_mem_ (red). The normalized fluorescence intensities on the dotted lines are shown on the right. **b**, Schematic representation of cpGFP insertion into the IL3 of OTR. **c**, Sequence alignment of the regions of the IL3 of human dopamine receptor D1 (hDRD1), a scaffold of a dopamine sensor (dLight1), and meOTR. **d**, Representative images of HEK293T cells co-expressing the indicated meOTR-cpGFP chimera (green) and mSca_mem_ (red). **e**, Pearson correlation coefficients comparing the meOTR-cpGFP chimeras and mSca_mem_ are summarized as the mean ± SEM (*n* = 13, 12, 12, 13, 12, 13, 13, 11, 13, 13, 13, 12, 13, and 12 cells; left to right). Statistics: one-way ANOVA (*F*_13,167_ = 1.78, *P* = 5.5×10^−32^) with Bonferroni *post-hoc* test (*P* = 5.1×10^−9^: N236–S269, *P* = 7.9×10^−14^: N236–I268, *P* = 1.2×10^−13^: F237–S269, *P* = 4.1×10^−14^: F237–I268, *P* = 3.1×10^−12^: K238–S269, *P* = 4.9×10^−14^: K238–I268, *P* = 1.6×10^−16^: L239–S269, *P* = 2.9×10^−14^: L239–I268, *P* = 1.2×10^−9^: K240–S269, *P* = 9.6×10^−11^: T241– S269, *P* = 5.4×10^−15^: T241–I268, *P* = 2.7×10^−16^: K242–S269, *P* = 2.0×10−^16^: K242–I268, compared with K240–I268). \*\*\**P* < 0.001. **f**, Summary of the insertion site-dependent characteristics of meOTR-cpGFP chimeras. **g**, Traces showing the fluorescence responses of the K240–I268 meOTR-cpGFP chimera upon stimulation with 100 nM OT. Scale bars, 10 μm (**a**, **d**).

**Extend Data Figure 2.**
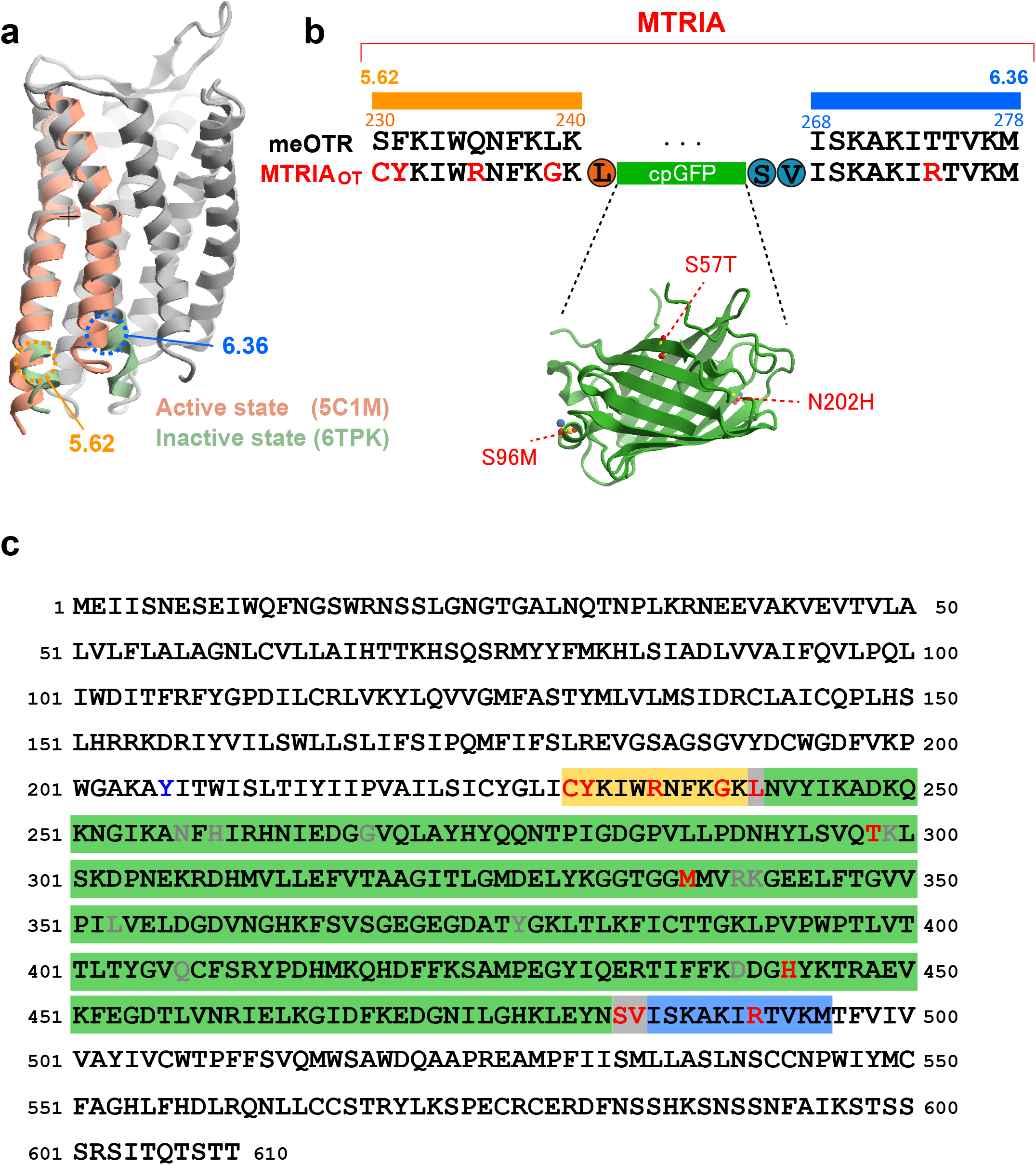
Sequence of a new fluorescent OT sensor. **a**, Superposition of active- and inactive-GPCR structures, adapted from the Protein Data Bank (PDB) archives (IDs: 5C1M and 6TPK). TM5–TM6 regions of the active and inactive states are colored dark salmon and dark sea-green, respectively. **b**, Alignment of the TM5–TM6 region between meOTR and MTRIA_OT_; the structure of cpGFP is adapted from a PDB archive (ID: 3SG2). The mutations introduced in MTRIA_OT_ are shown as red. **c**, Full amino acid sequence of MTRIA_OT_. The mutations in MTRIA_OT_, the point mutation in MTRIA_OT_-mut, and the mutations of cpGFP that were adapted in the other fluorescent sensors are shown in red, blue, and gray, respectively.

**Extend Data Figure 3.**
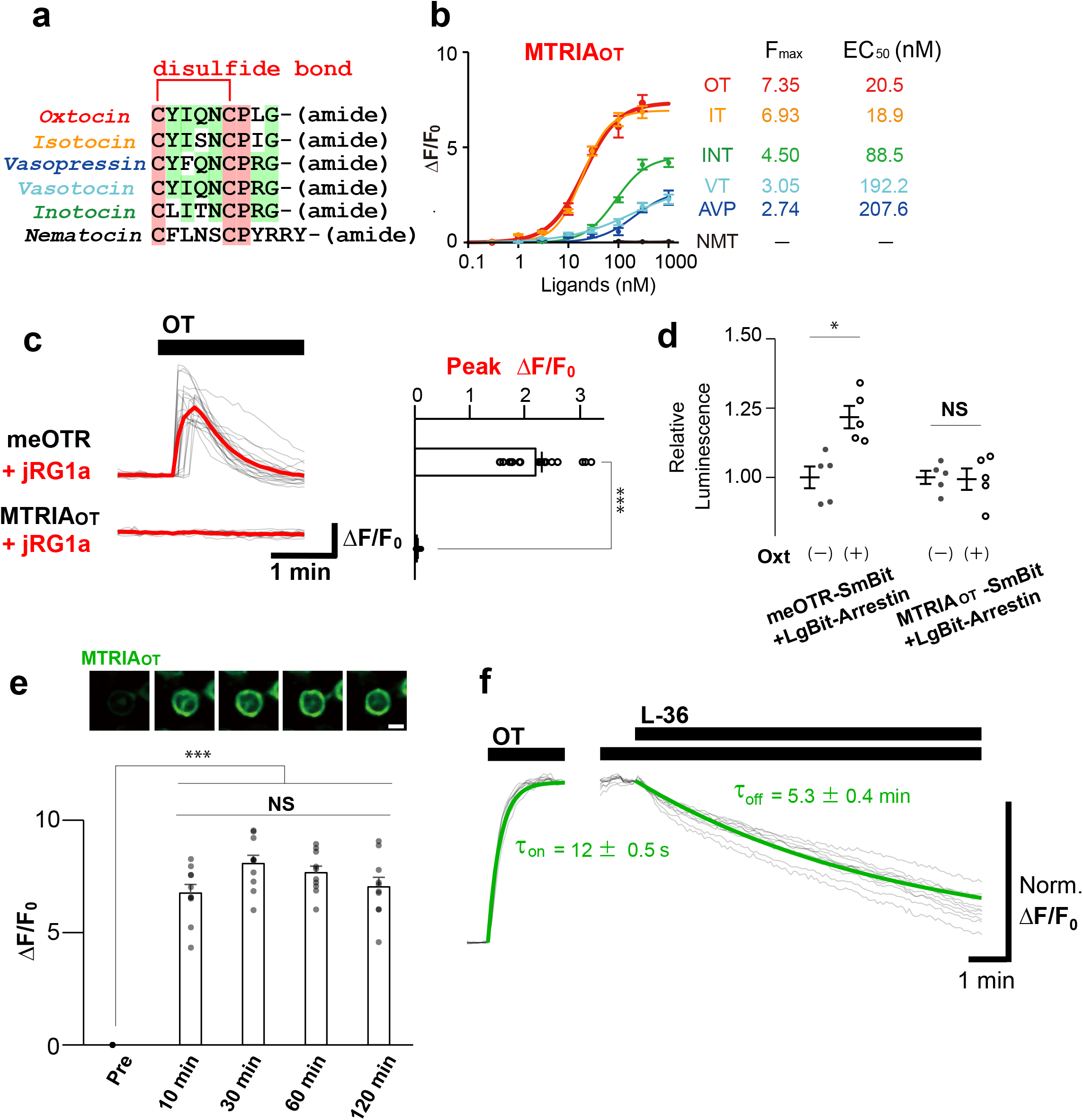
Basic properties of MTRIA_OT_. **a**, Sequence alignment of OT and its orthologous neuropeptides. **b**, Dose–response curves of MTRIA_OT_ to OT and its orthologous neuropeptides. **c**, Assay for G protein coupling. Traces of jRG1a fluorescence responses in either meOTR- or MTRIA_OT_-co-expressing cells upon stimulation with 100 nM OT (gray: each trace, green: average trace). Summary of the peak Δ*F*/*F*_0_ responses are shown on the right (*n* = 20 cells). Statistics: unpaired two-tailed *t*-test (*P* = 8.1×10^−21^). **d**, Assay for β-arrestin coupling. Traces showing the relative luminescence increase induced by NanoLuc luciferase complementation upon stimulation with 100 nM OT (*n* = 5 wells). Statistics: one-way ANOVA (*F*_3,16_ = 3.24, *P* = 0.0009) with Bonferroni post-hoc test (*P* = 0.029: (−) vs. (+) in meOTR-SmBit + LgBit-arrestin, *P* = 1: (−) vs. (+) in MTRIA_OT_-SmBit + LgBit-arrestin). **e**, Time-course of the fluorescence intensity of MTRIA_OT_ over 120 min following 100 nM OT stimulation. Representative images (top) and the summary of Δ*F*/*F*_0_ responses (bottom: *n* = 10 cells). Statistics: one-way ANOVA (*F*_4, 45_ = 2.58, *P* = 7.0×10^−22^) with Bonferroni post-hoc test (*P* = 9.7×10^−13^ Pre vs. 10 min, *P* = 1.9×10^−13^ Pre vs. 30 min, *P* = 9.3×10^−15^ Pre vs. 60 min, *P* = 2.6×10^−11^ Pre vs. 120 min, *P* = 0.24: 10 min vs. 30 min, *P* = 0.78: 10 min vs. 60 min, *P* = 1: 10 min vs. 120 min, *P* = 1: 30 min vs. 60 min, *P* = 1: 30 min vs. 120 min, *P* = 1: 60 min vs. 120 min). **f**, Traces showing the kinetics of MTRIA_OT_ in the fluorescence rise (left: *τ*_on_) and decay (right: *τ*_off_) upon OT binding and dissociation in HEK293T cells (gray: each trace, green: average trace; *n* = 10 cells). Scale bars, 10 μm (e). Graphs represent the mean ± SEM (**b**−**d**). \*\*\**P* < 0.001, \**P* < 0.05, NS: not significant (**c**−**e**).

**Extend Data Figure 4.**
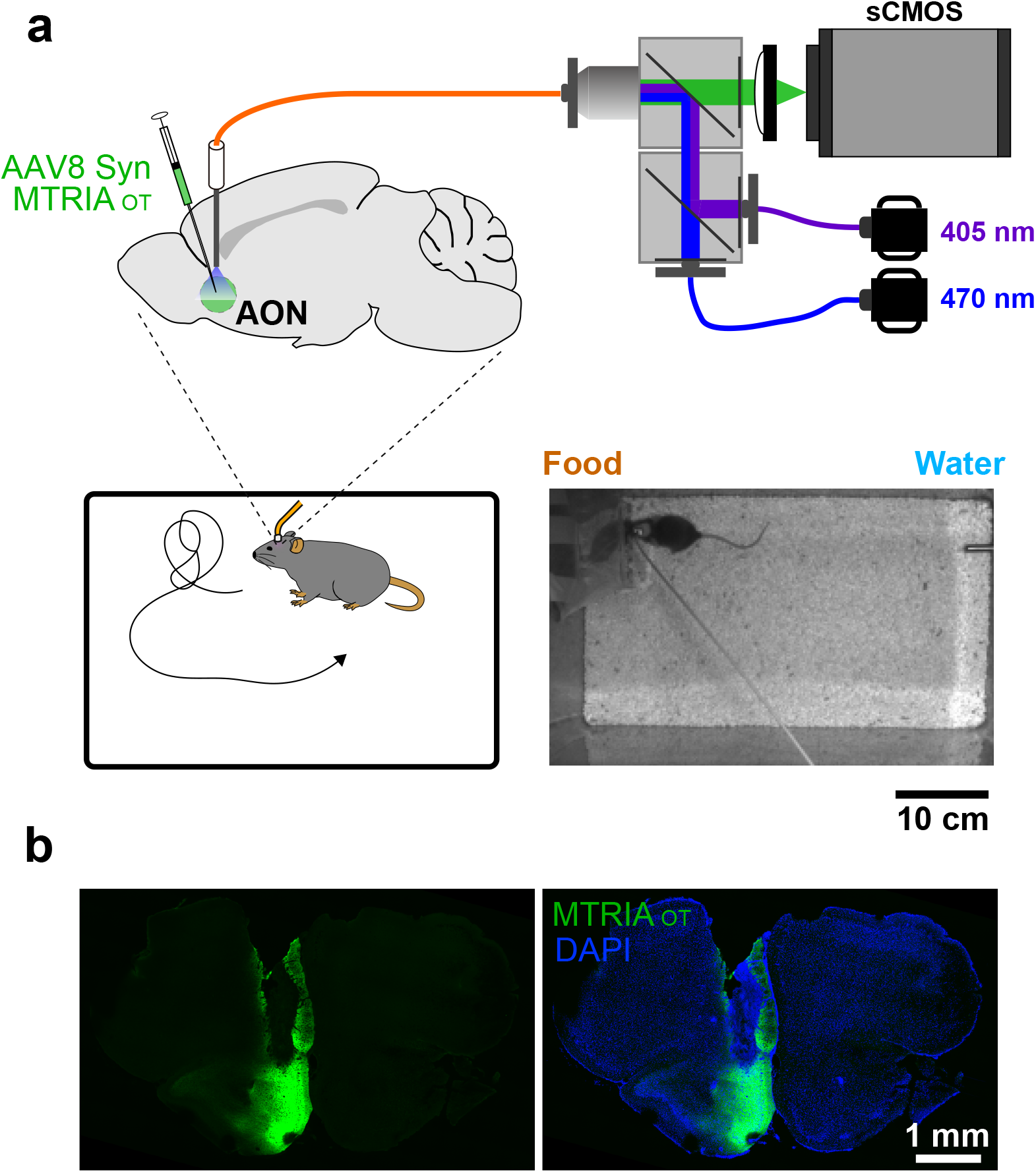
Fiber photometry setup and confirmation of the measurement site. **a**, Schematic illustrating the fiber photometry measurement of a freely behaving mouse. A representative image of an experimental arena captured by an overhead camera is shown in the bottom right. **b**, Histology showing the expression of MTRIA_OT_ (green) and the placement of an implanted cannula. DAPI (blue) was used for counterstaining nuclei.

**Extend Data Figure 5.**
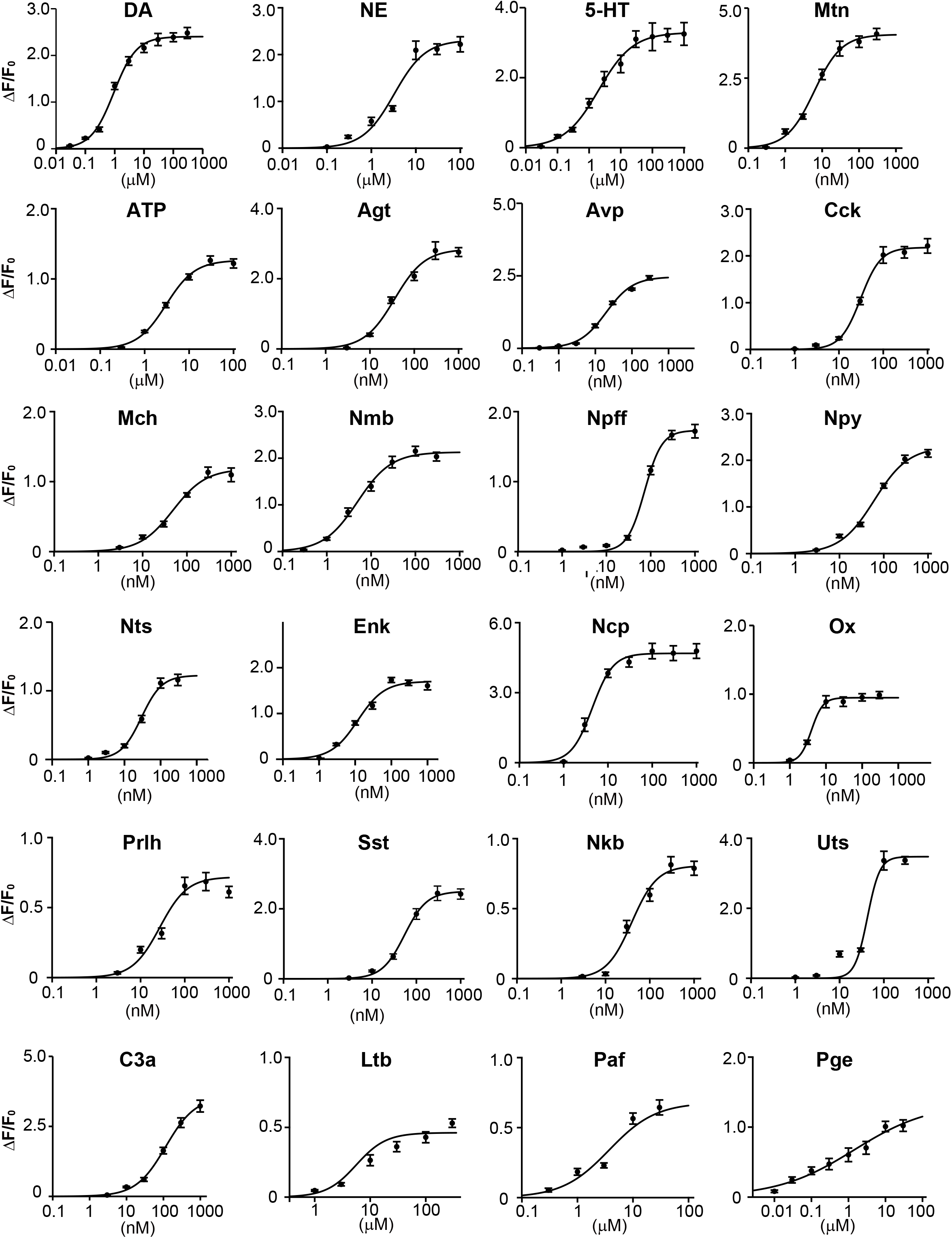
Dose–response curves of MTRIA sensors. Dose–response curves of the fluorescent sensors for 24 ligands that were examined in HEK293T cells. Data are shown as the mean ± SEM (*n* = 10 cells for each data point). The ligands were as follows: dopamine (DA), norepinephrine (NE), serotonin (5-HT), melatonin (Mtn), adenosine 5′-triphosphate (ATP), angiotensin (Agt), arginine-vasopressin (Avp), cholecystokinin (Cck), melanin-concentrating hormone (Mch), neuromedin B (Nmb), neuropeptide FF (Npff), neuropeptide Y (Npy), neurotensin (Nts), enkephalin (Enk), nociception (Ncp), Orexin (Ox), prolactin-releasing hormone (Prlh), somatostatin (Sst), neurokinin B (Nkb), urotensin (Uts), complement component C3a (C3a), leukotriene B (Ltb), platelet-activating factor (Paf), and prostagrandin E (Pge).

**Extended Data Table 1:**
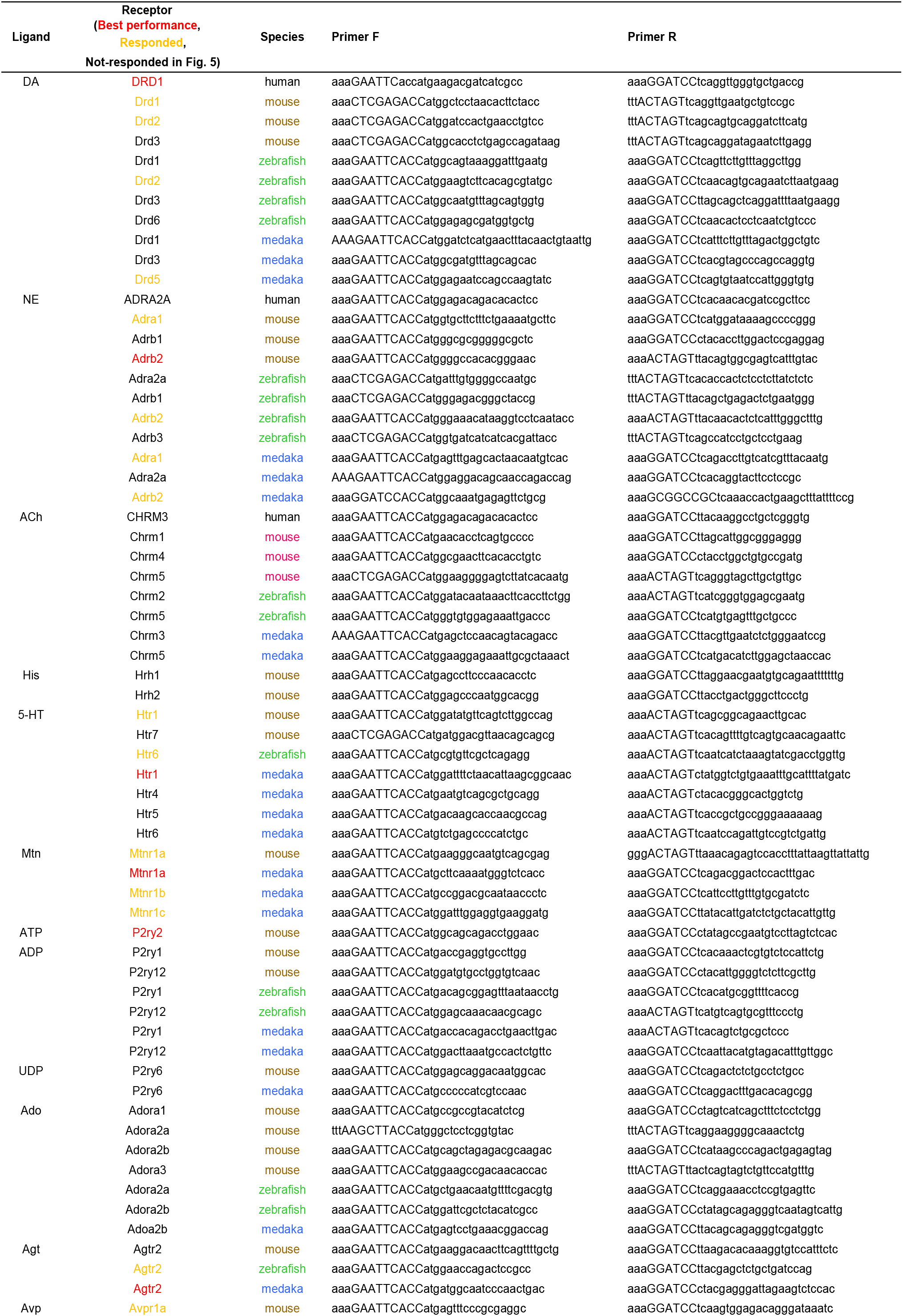
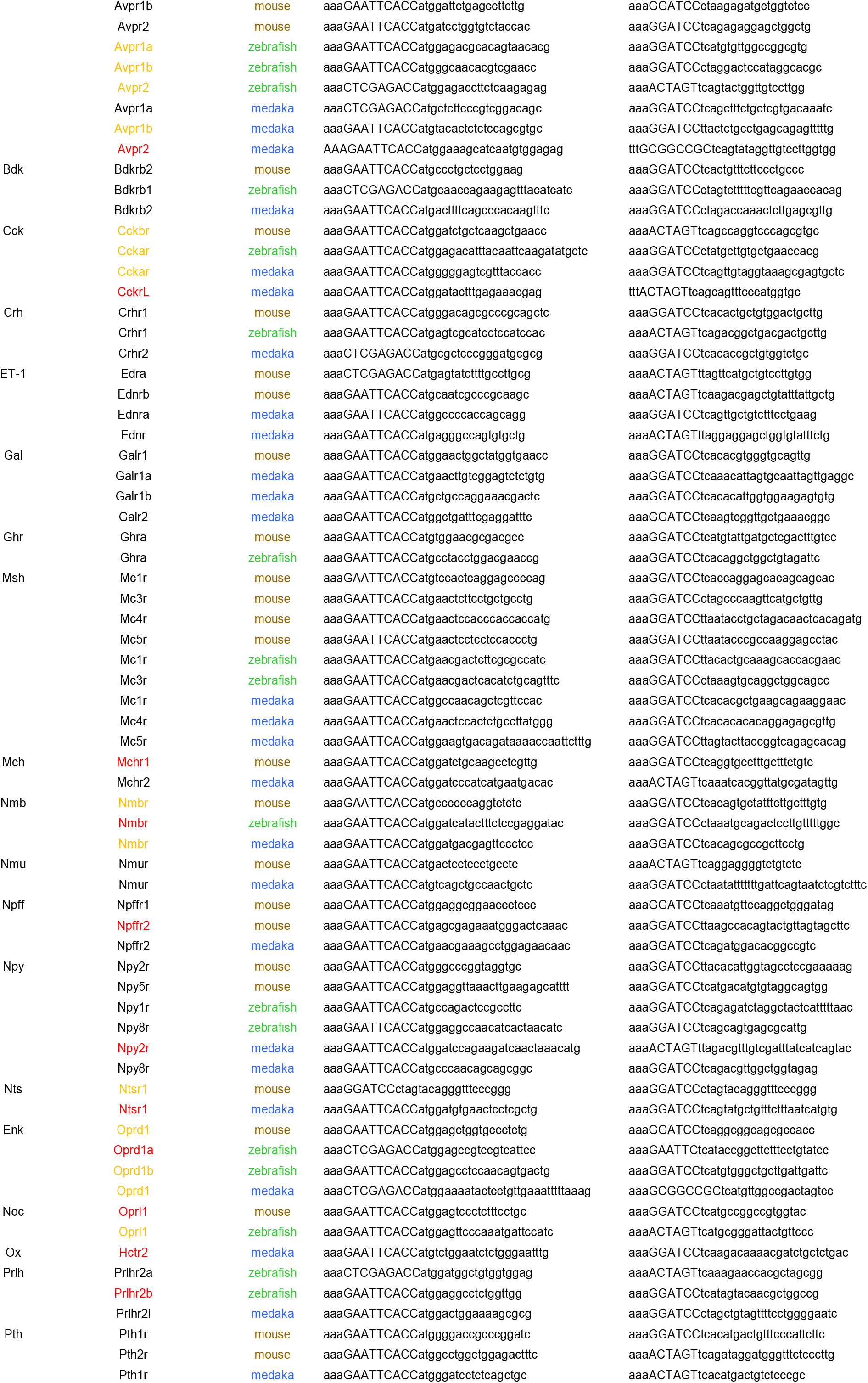
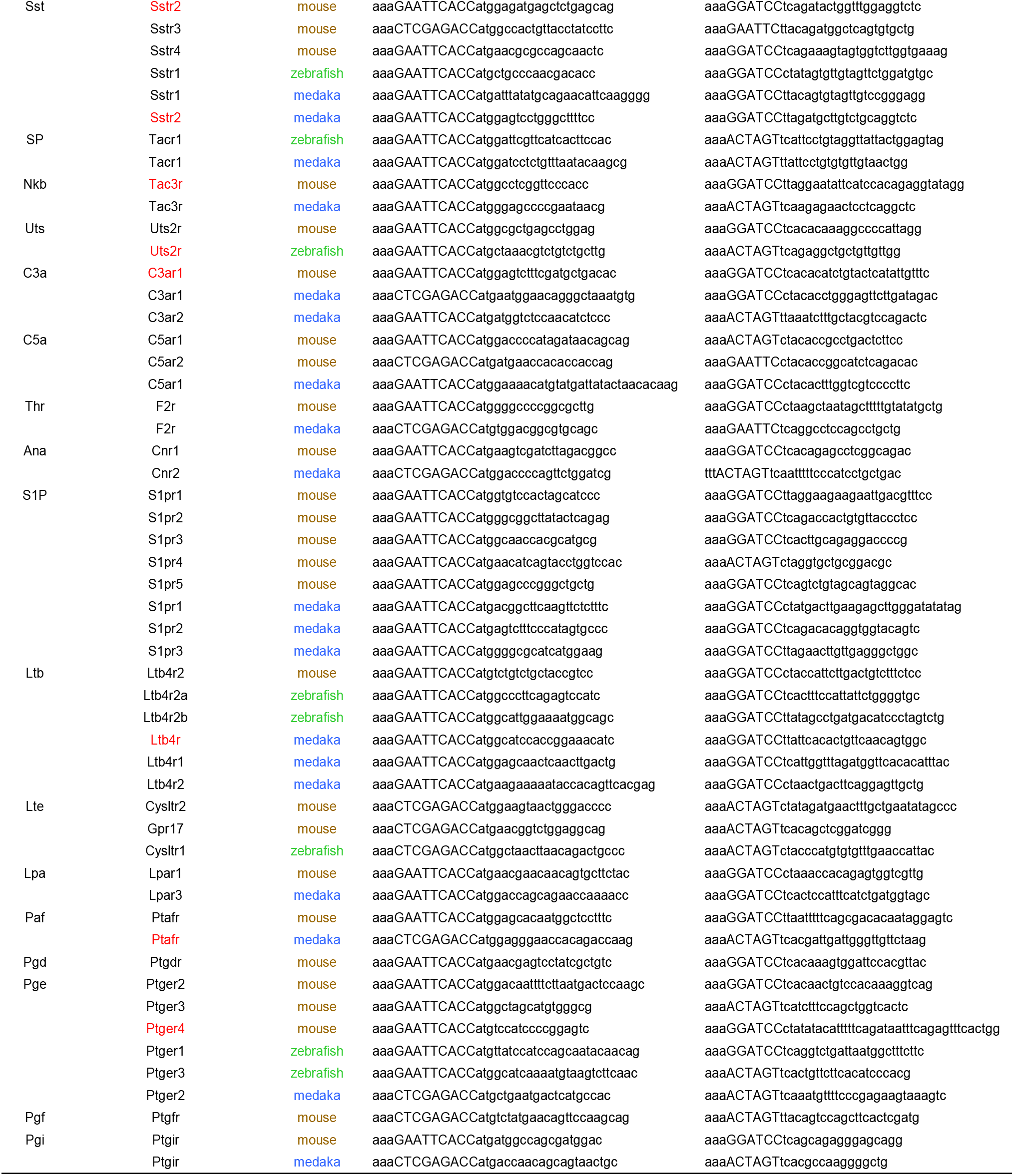
GPCRs used in Fig. 5 and the primer pairs for their cloning

## Notes

### Competing Interest Statement

The authors have declared no competing interest.

